# The ontogeny of play in a highly cooperative monkey, the common marmoset

**DOI:** 10.1101/2024.05.28.595935

**Authors:** Alice M. Godard, Judith M. Burkart, Rahel K. Brügger

## Abstract

Play is mostly observed in juveniles in mammals, and the type of play (social, locomotor and object play) tends to mirror adult function. In some species, also adults play with immatures, in particular if same-aged play partners are lacking and adults also invest in caretaking. We studied the ontogeny of play in cooperatively breeding common marmoset groups composed of parents and twin offspring between the age of two to six months. Social play was by far the most prevalent and increased with age. Adults were important play partners: Before 19 weeks old, the play partners of immatures was an adult in 54% of the time spent playing socially. After week 19, this proportion decreased to 29%. The rest of the social play time was spent playing with their twin. Thus, despite the constant presence of a twin, adult-immature play remained considerable, with equal contributions by mothers and fathers and no trade-offs with other care-taking behaviours (i.e., carrying and food sharing) for either of the parents. Notably, parents avoided playing simultaneously, presumably to avoid periods when no one could be vigilant. Together, these results resonate strongly with the highly interdependent and cooperative lifestyle of common marmosets.

**Highlights:**

- Social play is the predominant form of play in marmosets and increases between the age of 2 to 6 months
- Parents are important play partners for immatures during ontogeny
- The odds of playing with the twin rather than a parent increases with age
- Mothers and fathers play at similar amounts with the immatures
- Mothers and fathers take turns and almost never play simultaneously with the immatures

## Introduction

Play behaviour is widespread among mammals, and some orders are especially known for their playfulness, such as rodents, carnivores or primates (Burghardt, 2005; 2014). In all those orders, play is most prominent in juveniles, even though it can sometimes persist into adulthood as observed in dogs *Canis lupus familiaris*, bonobos *Pan paniscus*, or humans (Boere et al., 2020; Bradshaw, 2015; Burghardt, 2005; Hare et al., 2007; Pellegrini, 2009).

In the non-human animal (henceforth animal) literature, play is generally split into object play, locomotor-rotational (L/R) play and social play (Burghardt, 2005). Object play involves object manipulation without any apparent purpose, but is not always easily distinguished from object exploration and tool use (Pellegrini & Smith, 2005). L/R play is usually defined as any locomotor movement e.g., jumping, running, rolling, again with the additional specification of having no apparent purpose. Object and L/R play can either be solitary or integrated into social play. Social play involves two or more players that are usually conspecifics (Graham & Burghardt, 2010). The latter is the most frequently studied of the three play types and is also the most prominent in mammals and birds (Burghardt, 2005). Within social play, play fighting also known as rough-and-tumble play is most recognizable. Rough-and-tumble play mimics real fights and aggressive behaviours but the players very rarely come out of a play fight wounded (in humans: Smith, 1997), conversely to real aggressive contexts (Palagi et al., 2016; Pellis & Pellis, 2017). Interestingly, due to its quite intense nature, most often rough-and-tumble play involves only two partners and is rarely polyadic. The higher the number of partners, the harder it gets to negotiate the terms and make sure it is ‘only’ play and could hence be considered a feature of more tolerant species (Palagi and Cordoni 2012). Play does not occur uniformly throughout an animals’ life but follows an inverted-U-shape distribution with an early appearance during infancy, a peak at juvenile age and a slow decrease after this period (Panksepp, 1981; Pellegrini & Bjorklund, 2004). The fact that play behaviour almost disappears at a certain stage of an individual’s life indicates either that benefits from playing can only be gained during the immature period (Pellis et al., 2010) or that the costs are too high at a certain point (Pellis & Iwaniuk, 2000). Three main costs associated with play behaviour have been identified in the literature: increased predation risks (Harcourt, 1991; Hausfater, 1976), energy expenditure (Kunz et al., 2024; Pellegrini et al., 1998) and risks of accidental and social injuries (De Oliveira et al., 2003). Unfortunately, only a few studies have directly investigated these risks and quantified the costs during play e.g., predated animals during play, energy loss in kilocalories, wounds after a play fight (Pellegrini et al., 1998).

Despite the associated costs, immatures are highly motivated to play. Human children who have been stripped from their opportunities to play for some period, show increased play durations when opportunities can be resumed (Pellegrini et al., 1995; Pellegrini & Davis, 1993; Smith & Hagan, 1980). The same has been shown for social play in multiple species, such as wistar rats *Rattus norvegicus domestica* (Holloway & Suter, 2004), calves *Bos taurus* (Jensen & Kyhn, 2000 but see Bertelsen & Jensen, 2019) or pigs *Sus scrofa* (Wood-Gush & Vestergaard, 1991). Moreover, social play deprivation leads to less flexibility in animals’ social behaviour and impaired social reactions (e.g., Syrian hamsters, *Mesocricetus auratus*: Cooper et al., 2023; Wistar rats: Hol et al., 1999). This combination of high urgency to play and costs of playing in immatures suggests that there is an important biological function to this behaviour (Bond & Diamond, 2003).

There is no strict consensus on the ultimate function of play in the literature. One dominant view has been that play is a way for immatures to acquire skills for their adult life (Bateson, 2005; Pellegrini, 2009). Those deferred benefits encompass hypotheses about a motor training function (Berghänel et al., 2015 but see Byers & Walker, 1995; Pellis et al., 2023), an enhancement of social skills (Ahloy Dallaire & Mason, 2017; Blumstein et al., 2013; Pellis et al., 2010 but see Sharpe, 2005), a cognitive training function (Biben et al., 1989) as well as ‘training for the unexpected’ (Spinka et al., 2001). For example, object play in immatures has been especially studied for its potential role in acquiring skills to use tools in adulthood. Immature chimpanzees (*Pan troglodytes*), where adults are known to be skilled tool users, engage in object play, whereas their sister taxa bonobos, who are not frequently using tools, show comparatively less object play (Koops et al., 2015; Pellis & Pellis, 2017; Pellegrini & Smith, 1998, but more informations are needed to conclude on the developmental trajectory of object play in these two species, see Koops, 2024). Similar findings exist in several avian species, where the type and frequency of object manipulation was dependent upon the species ecology (Auersperg et al., 2015). Even at the individual level in the same species, the way an individual plays with objects seems to have an effect on the expression of tool use later on, such as in longtailed macaques *Macaca fascicularis* (Cenni et al. 2024).

Play is clearly the realm of juveniles, but in certain species adults continue to play at a high rate, for example in humans (Proyer 2018; Van Vleet & Feeney, 2015), bonobos (Palagi, 2023) or African wild dogs *Lycaon pictus* (Spiotta, 2022; Frame et al., 1979). This is puzzling because the important benefits of play are mainly associated with the juvenile period and the same costs described for the immatures apply to the adults as well and should prevent them from playing. Adults do play at the expense of their caloric intake and time. They also compromise their own and offspring safety by lowering their vigilance (Harcourt, 1991; Hausfater, 1976). Indeed, while immatures are playing, they are more vulnerable to predators because they are more conspicuous. Adults would thus be expected to protect the immatures with increased vigilance instead of playing themselves and risking of being surprised by a predator (Biben et al., 1989; Harcourt, 1991; Hausfater, 1976), as it has already been demonstrated In golden lion tamarins *Leontopithecus rosalia* (De Oliveira et al., 2003).

Certain factors seem to favour play behaviour of adults. First, the level of social tolerance in a group (Palagi, 2023; Palagi, Cordoni, et al., 2016) could facilitate adult-immature play. Further, the level of investment in infant caretaking displayed by a parent or an adult of the group might be an indicator as to whether this parent or adult will spend time playing with the infant. In other words, a parent or adult that spends time with the infant, and takes care of it, is more likely to play with the infant than a random individual of the group e.g., in chimpanzees *Pan troglodytes* (Pellegrini & Smith, 2005), orangutans *Pongo spp.* (Fröhlich et al., 2020; Kunz et al., 2024) or wolves *Canis lupus* (Essler et al., 2016) although this is not always the case. For example, in squirrel monkeys *Saimiri boliviensis* (Biben et al., 1989) and rhesus macaques *Macaca mulatta* (Kulik et al., 2015; Tartabini, 1991), even mother-infant play is rarely or never observed. A factor that could explain these differences between species is the presence or absence of other immatures. It has been suggested that immatures prefer playing with their peers, and will play with adults if there are no other options (Kunz et al., 2024; Pellis & Iwaniuk, 2000; Poirier, 1970). Another dimension of parent-immature play is the age of the immatures, play with parents being more prevalent in young infants than juveniles (Amodia-Bidakowska et al., 2020; Kunz et al., 2024; Mackey et al., 2014; Paquette & StGeorge, 2023). Ultimately it has been suggested that, by playing with immatures, adults facilitate the immatures’ acquisition of species-specific signals and skills employed in various social interactions (Enomoto, 1990; Mackey et al., 2014; Paquette & StGeorge, 2023; Sutton-Smith, 1993). Adults would thus invest time to pass down, voluntarily or not, some social skills (Paquette et al., 2003) and may ultimately contribute to increasing the fitness of the immatures (Amodia-Bidakowska et al., 2020). However, there is still no clear answer as to why parents having an active role in caretaking do not always play with their offspring.

The goal of this study was to investigate the ontogeny of play, with a special focus on parent – immature play in common marmosets. Marmosets are cooperative breeders, where all group members provide care for the immatures and are part of the broader family of callitrichids. Callitrichid monkeys are known for their high level of social tolerance and general prosociality, characteristics that are directly linked to their cooperative breeding system (Burkart & van Schaik, 2020; Hrdy & Burkart, 2022; Schaffner & Cain, 2000). Consequently, their high social tolerance and prosociality are often studied through the lens of cooperative acts performed to take care of the immatures, mainly with two specific behaviours: infant carrying and provisioning the immatures with food, in captivity as well as in the wild (Burkart and van Schaik, 2020; de Oliveira Terceiro et al., 2021). Groups are usually composed of two breeders and several sexually mature helpers. Adult breeders and helpers appear to show a genuine concern about the immatures’ well-being (Brügger et al., 2018, 2023) and proactively share food and information (Sehner et al., 2023). Males are important caretakers for the immatures, in contrast to most of the other primate species (Erb & Porter, 2017). Additionally, females generally give birth to twins (Haig, 1999). Female breeders experience high reproductive costs due to twinning and post-partum oestrus (Beehner & Lu, 2013; Leutenegger, 1973); male breeders and helpers are therefore crucial to alleviate the energetic costs on the females during the first months of the immatures’ life. Here, we longitudinally documented play, in three marmoset groups with twin immatures between two and six months old. All groups were composed of two breeders and two immatures without helpers.

In part 1 (Play trajectories), we investigated the ontogeny of object, L/R and social play, predicting that all three categories should increase in frequency as the immatures grew up, but that social play should rapidly become the most important category of play. Until one month old, marmosets are not independent and are carried by their parents or helpers (Abbott et al., 2003). They can still be carried between one and two months as well, more or less frequently and they are still learning how to walk during this period (Schultz-Darken et al., 2016). We therefore expected solitary play to be more important early in development and peak before social play, the latter requiring more accurate control over one’s own body and others’ understanding.

In part 2 (Play partners), we investigated immatures partners. In early studies on common marmosets (Voland, 1977) it has been found that immatures prefer to play with their twin rather than adults, but Stevenson & Poole (1982) suggested that partner preference depends on the age of the immatures and adults could be important play partners at an early stage of the immatures’ development. This pattern has also been found in several other species (Amodia-Bidakowska et al., 2020; Kunz et al., 2024; Mackey et al., 2014; Paquette & StGeorge, 2023). Knowing that marmosets are highly tolerant and adults contribute actively to immature care, we thus predicted that adults would engage in play at high rates with the immatures, especially at early developmental stages, in dyadic as well as in polyadic play. We nonetheless expected triadic play to be less present than dyadic play, as it is harder to maintain.

In Part 3 (Parental investment and coordination), zooming in on parent-immature play, we made three predictions. First, we looked at each parent separately, to see if mothers and fathers would play to a similar extent with the immatures. Stevenson & Poole (1982) found that, in common marmosets, adult males played more than did adult females, and they showed that it was linked to the reproductive state of the females. It is likely that they have to compromise by playing less, since it is an energy-consuming behaviour (Guerreiro Martins et al., 2019). In their study, Stevenson & Poole (1982) observed play in extended family groups comprising helpers but since no helpers were present in our study groups, we expected that the difference between fathers and mothers’ playful investment would be even higher, with male breeders being much more playful and with female breeders having less energy to allocate to playing in the absence of helpers. Second, we considered commonly described caretaking behaviours, carrying and food provisioning, to see if they would positively correlate with playing, regardless of the sex of the parent. Since the level of investment in infant caretaking displayed by a parent might be an indicator as to whether this parent will spend time playing with the immature, we had two competing hypotheses. On the one hand, parents investing time and energy in infant rearing would be expected to invest more in play behaviours with the immatures than less involved parents. On the other hand, it could be that there is a trade-off in parents between caretaking behaviours and play. We would therefore observe a negative correlation between play and carrying as well as food-sharing. Third, since play can take a toll on the immatures’ fitness by reducing their focus on antipredator vigilance, we expected parents to take into account this risk and act accordingly by taking turns when playing, especially in small groups. Callitrichids are indeed known to coordinate vigilance. For instance, marmoset dyads take turns in being vigilant and feeding (Brügger et al., 2023; Phaniraj et al., 2023). When a parent plays with immatures, it can not be vigilant at the same time. The only solution to stay vigilant at the group-level is therefore that the other adult stays vigilant instead of joining the play bout. We thus examined whether and how much the father and the mother played simultaneously i.e., we searched for all overlaps of time when both parents were playing to see if they actively avoided playing together with the immatures or not.

## Methods

### Subjects and housing

We collected observational data on three family groups (N = 12) of captive common marmosets. The groups were observed when immatures were between 2 and 6 months old (see additional information for group composition and data collection details in Table S1). All groups had immature twins and none had helpers (Abbott et al., 2003). The groups were thus composed of two breeders and two immatures.

All individuals were captive-born, parents-reared and housed in family groups. Their heated indoor enclosures were equipped with various climbing materials (branches, ropes, tubes, and platforms), a sleeping box, an infrared lamp, and floors covered by bark mulch. The animals had regular access to outdoor enclosures when weather conditions were adequate. Feeding of all animals occurred once in the morning at 08h00 with a vitamin-enriched mash and at around 11h30 with a variety of vegetables. During the afternoon, the marmosets received diverse additional protein sources such as insects or eggs. Before, during and after the observations, water was always available through water dispensers.

### Data collection

#### Video-recordings for play data

All sessions were collected with either two or three GoPro cameras (Hero7 white with a resolution 1920*1440 pixels and Hero9 black with a Wide Quad HD resolution). The cameras were placed to ensure almost full visual coverage of the enclosure by placing one camera at the front, one at the back of the enclosure, and additionally one on the ceiling, for a duration of 30 minutes. The trap door leading to the outdoor enclosures was always closed during the recordings. No experimenter was present during the video-recordings.

To provide an attractive play platform, a hammock (large towel) was placed in the enclosure and clipped to the mesh ceiling during the sessions. Old jeans or towels are already part of the standard equipment of home enclosures and marmosets spend a lot of time playing in/on them. We thus replaced them by a new hammock (large towel) in a way that allowed optimal visibility for the cameras.

We collected data on the three groups regularly between 2 and 6 months of the age of the immatures, keeping the balance between the different periods of the day to avoid any unwanted bias (Norscia & Palagi, 2011): before the main feeding (BF), 30 minute videos-recordings within the morning period 09h–11h and after the main feeding (AF), 30 minute videos-recordings within the afternoon period 13h–14h. Two sessions (one BF and one AF during the same week) were recorded almost every other week for each group during five months. Overall, a total of 66 sessions were collected corresponding to 1980 minutes of video material. All the individuals from the groups were visible during the recordings, which means that 660 minutes of videos were collected for each individual of each group.

#### Food sharing and carrying data

For each breeder, immature carrying was recorded daily between 8h and 17h in hourly group scans for 120 days after birth. This means that, for each breeder, the number of infants carried (either 1, 2 or 0) was recorded every hour during the day. To evaluate how much food each parent would share with the immatures depending on their age, we followed the food sharing protocol described in Guerreiro Martins et al. (2019), twice a week during the whole data collection period (see Table S1). The high-value food to be shared, a defrosted cricket, was distributed in a standardised manner, one item at a time and one adult at a time. During each food sharing session three crickets, i.e., three trials, of approximately the same size were sequentially provided to each breeder, and not at the same time, meaning that in total there were six trials per food sharing session. For each trial, we observed if the cricket was shared or not and if it was shared proactively (food calls from the donor before any begging from the immatures), reactively (begging from the receiver happened before the donor food called if she or he food called at all), resisted (food was transferred but adults showed resistance by chatter calling and/or running away from the begging individual), and who the recipient was. If the cricket was not shared, we also recorded if others had begged (refusal, i.e., no sharing despite begging). Therefore, for each trial, the recipient(s) of the cricket could be: a) none, b) either one of the immatures, c) both of the immatures or d) both of the infants and the other breeder. During the first months we could not distinguish between the twins as they had no distinguishable marks; they were thus counted as “immatures”. AMG and RKB collected the food sharing data for the Washington and Jambi groups. Sandro Sehner collected data for the Guapa group. We ensured inter-observer reliability by having 14% of all food sharing trials collected by two raters simultaneously. A Cohen’s kappa of 0,64 was reached.

#### Ethical Note

All experiments were carried out in accordance with the Swiss legislation and licensed by the Kantonales Veterinäramt Zürich (licence number: ZH232/19). All of the experimental procedures were fully non-invasive (degree of severity: 0) as most of them were only behavioural observations and the rest (food-sharing experiments) were about giving the individual highly valued food items (crickets). No experiments were conducted outside the home enclosure and no animal was food or water deprived during any point of this study.

### Data coding for play data

We conducted frame-by-frame analyses (25fps or 29.97fps) using the software INTERACT (Mangold GmbH, version 18.7.7.17). We coded the middle 15 minutes (07m30s–22m30s) or 5 minutes (12m30s–17m30s) from the videos that were used, using focal sampling (Altmann, 1974). All individuals were present and coded in each video analysed and although the immatures were not individually identified because they were too young, we were able to always follow both immatures separately and code them both independently, removing any potential bias. In total across the three groups, we coded 1060 minutes. This represents approximately 20 minutes of video footage in one or two different sessions for each month of the immatures’ life between 2 and 6 months and for all individuals of each group.

### Play definition

We identified play behaviour following Burghardt’s (2005) five criteria:

1. The behaviour is not fully functional in how it was expressed, and thus did not contribute to immediate survival
2. The behaviour is spontaneous, voluntary, intentional, pleasurable, rewarding, reinforcing, or autotelic
3. Compared to functional behaviours it is either incomplete, exaggerated, awkward, precocious, or modified
4. The behaviour is repeated throughout the development of the individuals, but not in a rigid stereotypical way
5. The behaviour is observed when the animals are relaxed and not under stress or competition

Further, we made sure to exclude real fight situations from social play. During real fights both the aggressor and the victim produce chatter call vocalisations, and the attacks are only unilateral with a clear dominant (Bezerra & Souto, 2008; Stevenson & Poole, 1976). The bites are also usually not inhibited which means that wounds can be visible afterwards. Moreover, intra-group aggression rates in marmosets are known to be rare (Finkenwirth & Burkart, 2018).

We defined play operationally as any social or solitary element present in our ethogram occurring while focal-sampling the individuals. Social play included two or more individuals and solitary play included object manipulation and locomotor/postures play without a play partner.

### Ethogram

The ethogram (Figure 1) was made using the marmoset play ethograms of Stevenson & Poole (1976, 1982) and Voland (1977). Further additions were made with the help of other play behaviours described in Petrů et al. (2009) and Cordoni & Palagi (2011). In addition to the illustrated ethogram, supplementary video material of all coded play behaviours is provided (Section S3).

**Figure 1.**
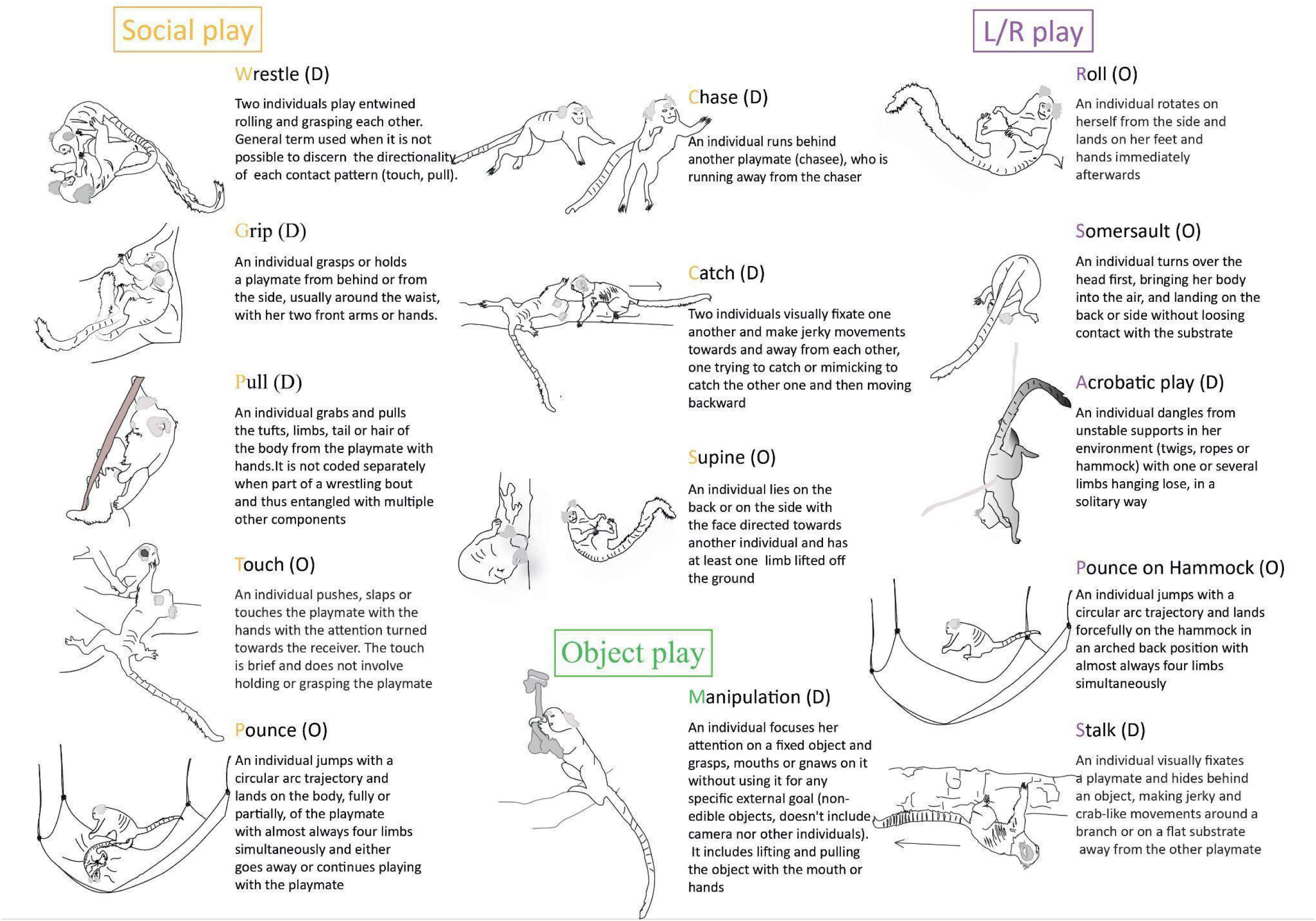
Illustrated ethogram with all the play elements coded in the present study. The playful elements were classified into three bigger categories: Social play, L/R play and object manipulation. (D): play element coded with a duration, (O): play element coded as an occurrence.

A coding protocol (see the protocol in Table S2) with refined definitions was established to facilitate recognition of each play element. For components with a duration, this included when the exact start and end had to be coded (see Figure 1 for details about which play elements were coded as durations or occurrences). Importantly, the following rules were implemented during video-coding:

1. For the social play elements, the focal individual was always chosen in function of the initiation: she was chosen if she was the first one to make physical contact i.e., during wrestle or pulling, or the one doing the action i.e., chasing another one or doing a supine.
2. No individual could be twice a focal individual at the same time. In other words, an individual could not be performing several actions simultaneously in our coding scheme. To ensure that only one action would be coded per individual at any time point, we decided to prioritise social elements before locomotory ones e.g., two individuals wrestling in an acrobatic position because they were hanging from the hammock was only coded as wrestling and not as doing an acrobatic posture. In solitary L/R play, by definition there is no partner thus it made sense to not code a solitary play element concomitant to a social play element as it would be a contradiction. However, if there was a sequence of social play elements followed by solitary play elements and then the social play elements were resumed, the solitary elements were counted as solitary elements and not omitted because they were not concomitant but in a sequence. We then prioritised elements with durations before occurrences e.g., if an individual chasing another one and managed to touch the chasee while running, the touch was not coded.

If one individual was totally hidden behind an object in the enclosure and thus not visible, she was coded as out of sight (oos). If two individuals were playfully interacting but the behaviour was either not described in the ethogram or not fully visible from the cameras, it was coded as ‘unknown’ (UK), thus acknowledging that there was play but without inferring what it was. UK play elements could either have duration or be coded as states and could be solitary or social. For all analyses regarding play amount between dyads or triads, the UK play elements were included. We proceeded the same for solitary play elements that were not described in the ethogram.

Because the identity of the twins of each group were not recognisable, they were coded indiscriminately as immatures. In summary, the following variables were coded: players identity (immatures, father and mother), frequency or duration of the play items, including UK elements, directionality of the social play elements (who is performing the action towards whom), for the element “pull” what was the target (tail, hair of the body, tufts or limbs) and for the element “acrobatic play” where it was happening (towel, rope, twig).

We assessed inter-observer reliability for each play element with a second rater (Melanie Meyer) who coded 9,4% of the videos, by looking at intraclass correlation coefficients (ICC 3) for each play element that we coded, either by using the duration or the occurrence of those elements. Roll and somersault were excluded from the IOR because only one element of each was present in the selected video bouts for the IOR. Overall, those two components were very rare and thus excluded from all analyses (Table S3).

## Data preparation

### Part 1: Play trajectories

We calculated the frequency at which immatures performed social, locomotor and object play elements as the number of times that the immatures performed a social, locomotor or object play element out of the total time observed in a specific session. Because the immatures were not recognisable and both coded as ‘immatures’, the calculated play frequency then represents both immatures for the three categories. All play instances were taken into consideration, no matter the play composition (dyads or triads). We adjusted for the time the immatures could have been focal or receiver of a play element but one of the potential players was out of sight by subtracting from the total time observed the time when at least one individual was out of sight.

### Part 2: Play partners

A social play sequence consisted of different play elements following one another with interruptions between these play elements. These interruptions corresponded to the cessation of the play elements (e.g., cessation of contact after wrestling or pulling, chasing stops) before resumption, or not. The moment of resumption corresponded to the moment when another play element started between the same two, or three, partners. To belong to the same play sequence, or play bout, the resumption had to happen within a certain time interval.

Different interval lengths are found in the literature (Hawley, 2016: 1 minute for mountain gorillas *Gorilla beringeiberingei*, Heesen et al., 2021: 2 minutes for bonobos, Stevenson & Poole, 1982: 5 seconds for common marmosets, Govindarajulu et al., 1993: no interval for vervet monkeys *Cercopithecus aethiops sabaeus*), but the most common one in the literature across species is 10 seconds (Cordoni et al., 2016; De Oliveira et al., 2003; Iki & Hasegawa, 2020; Mackey et al., 2014; R. Wright et al., 2018), which we also chose for our study. Additionally, if the play partners changed or if one other player joined, we considered that the play bout had ended, even if the play elements were less than 10 seconds apart. We considered the arrival of a new player as a new play bout. Besides following the published literature we additionally explored how changing the intervals between play elements would affect the resulting amount of time playing (see this additional analysis in section S1), which corroborated our decision to adopt a 10 second interval.

Based on these calculated play bouts we calculated the proportion of time a dyad or triad was observed playing out of the total time observed in a specific coded session. The dyads were specified as either Dyads_PARENT-IMMATURE_ (where we added all the Dyads_MOTHER-IMMATURE_ and Dyads_FATHER-IMMATURE_ together) or Dyads_TWINS_. The triads were specified as Triads_PARENT-TWINS_ (where we added all the Triads_MOTHER-TWINS_ and Triads_FATHER-TWINS_ together). We adjusted for the time a dyad or triad could have been playing but one of the individuals was out of sight by subtracting from the total time observed the time when at least one individual was out of sight.

The play compositions were calculated as follow:

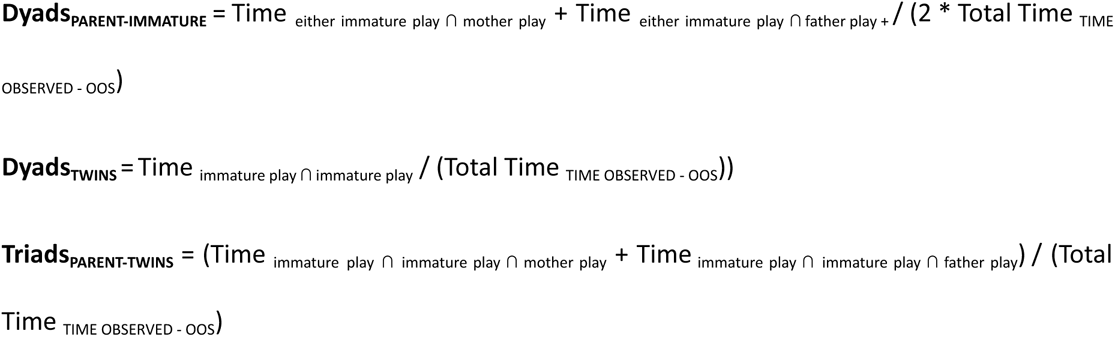

### Part 3: Parental investment and coordination

We compared female to male breeders’ investment in play by calculating the Play_MOTHER-IMMATURE_ and Play_FATHER-IMMATURE,_ as proportions of time spent playing respectively by the female breeder and the male breeder with one (dyad) or two immatures (triad), out of the total time observed in a specific session. We adjusted for the time a breeder or immature could have been playing but was out of sight by subtracting from the total time observed the time when at least one individual was out of sight.

We calculated the food sharing rate as the number of crickets reactively or proactively given over the total number of trials per month of the immatures’ life.

We calculated the carrying rate as the number of immatures carried i.e., we summed up the number of immatures, maximum 2, on every hourly scan, in total over the 120 days of carrying data collection controlling for the number of immatures the focal parent could have carried controlling for the times where no data was available i.e., missing data in the hourly scans.

We additionally determined the predicted probability of simultaneous play of both parents under the assumption that play bouts happen independently: P_MOTHER PLAY_× P_FATHER PLAY_ = Time_MOTHER PLAY_ / (Total Time _TIME OBSERVED-OOS_) × Time_FATHER PLAY_ / (Total Time _TIME OBSERVED-OOS_) and the observed probability: Time _Aplay ∩ Bplay_ / (Total Time _TIME OBSERVED - OOS_) for each session.

If the observed probability of simultaneous play is lower than the predicted probability, individuals take turns playing, if the opposite is true, individuals are more likely to play synchronously.

### Data analysis and statistics

All statistical analyses were conducted using the statistical software R studio (2023.12.1). We used a Generalized Linear model (GLM, package “stats”) and zero inflated Generalized Additive Models for Location, Scale and Shape (package ‘Gamlss’). Our outcome variable for model 1 is a discrete proportion (number of social, L/R or object elements over the total time observed in a specific session controlled for the time out of sight), we therefore chose to use a model from the poisson family. We controlled for overdispersion and adjusted accordingly by fitting a quasi-poisson. We also controlled for the absence of outliers (standardized residuals < 3), high leverage points (hat values < 3 * mean hat value) and influential cases (cook’s distance < 1).

For models 2, 3a and 3b, our outcomes are continuous proportions (time spent playing over total time observed controlled for time out of sight), we thus used a model from the zero-inflated beta family. For these models, worm plots were used.

The homogeneity of the residuals was assessed by inspecting residual plots for all models (function ‘plot’ and function ‘resid_plots’, package ‘gamlss.ggplots’). We controlled for the absence of collinearity between the predictors (function ‘vif’) for all models. The full models including all relevant predictor variables were always compared to the null models only including the intercept by using a likelihood ratio test (functions ‘anova’; package ‘car’ and function ‘lrtest’; package ‘lmtest’). Model selection was based on the Akaike Information Criterion AIC. Post hoc comparisons were conducted only for model 2 to discriminate between the groups (function “emmeans” and “emtrends”, package “emmeans”). We report the overall goodness-of fit of the models with conditional R2 values (function “r.squaredGLMM”, package “MuMIn” and function ‘Rsq’, package ‘gamlss’). Prediction plots were drawn using the functions ‘Effect’ (package ‘effects’), ‘ggplot’ (package ‘ggplot2’) and ‘predict_response’ (package ‘ggeffects’).

#### Part 1: Play trajectories

Our first analysis modelled the occurrence per coded session of play elements categorised as ‘social’, ‘object manipulation’ or ‘L/R’. The number of elements from each category was summed up per coded session (model 1). We controlled for the relative length of the coded session by setting the logarithm of the total observed time for the coded session in question, adjusted with the time out of sight as the offset. We included the play category (social, object, L/R), the age of the immatures (in weeks) and the group as well as all two-way interactions between play category and age of the immatures, as fixed effects. We set a planned contrast between the different categories of play, with the first contrast being social play against the two others categories and the second contrast comparing object versus L/R play.

#### Part 2: Play partners

Our second analysis allowed us to investigate the immatures’ partners. We calculated the proportion of dyadic and triadic play to compare parent-immature play to twin play as our outcome (model 2). We included the play composition (Dyads_PARENT-IMMATURE,_ Dyads _TWINS_, Triads_PARENT-TWINS_), the age of the immatures (in weeks) and the group as well as the two-way interaction between dyad identity and age as fixed effects. We set a planned contrast between the different play compositions, with the first contrast being dyadic (both Dyads_PARENT-IMMATURE_ and Dyads _TWINS_) against triadic play and the second contrast comparing the Dyads_PARENT-IMMATURE_ to the Dyads _TWINS_).

#### Part 3: Parental investment and coordination

To understand if mothers and fathers would play differently, we calculated the proportion of time they spent playing per coded session as our outcome and fitted a zero inflated Gamlss model (model 3a). Breeder sex (male, female), age of the immatures (in months) and group identity were used as fixed effects.

Additionally, we analysed whether there were other factors than breeder sex that could influence the involvement in play behaviour. To do so we ran another model (model 3b) with carrying rate, food sharing proportion, age of the immatures (in months) and group identity as fixed effects.

We then examined whether and how much the father and the mother played simultaneously. We used a paired Wilcoxon signed rank test (function ‘wilcox.test’, package stats) to compare the predicted probability of overlap to the observed probability.

## Results

### Part 1: Play trajectories

We first analysed the developmental trajectories of the three play categories (L/R play, object manipulation and social play). The full model (AIC = 988) that included the fixed effects play category, age of the immatures and group as well as the two-way interactions between age and play category, explained the data significantly better than the null model (N_total_ = 63, N_individuals_ = 6, N_groups_ = 3; AIC = 3278; likelihood ratio test: *X*^2^ (7) = 2303, *p* < 0.001). We found that immature marmosets did increase their amount of social play with age. There was a very low and homogeneous proportion of both object manipulation and L/R play throughout the 5 months of observations (Figure 2). Further, social play was the most prominent form of play all along the 5 months of observation (see Table 1 for model summary).

**Figure 2.**
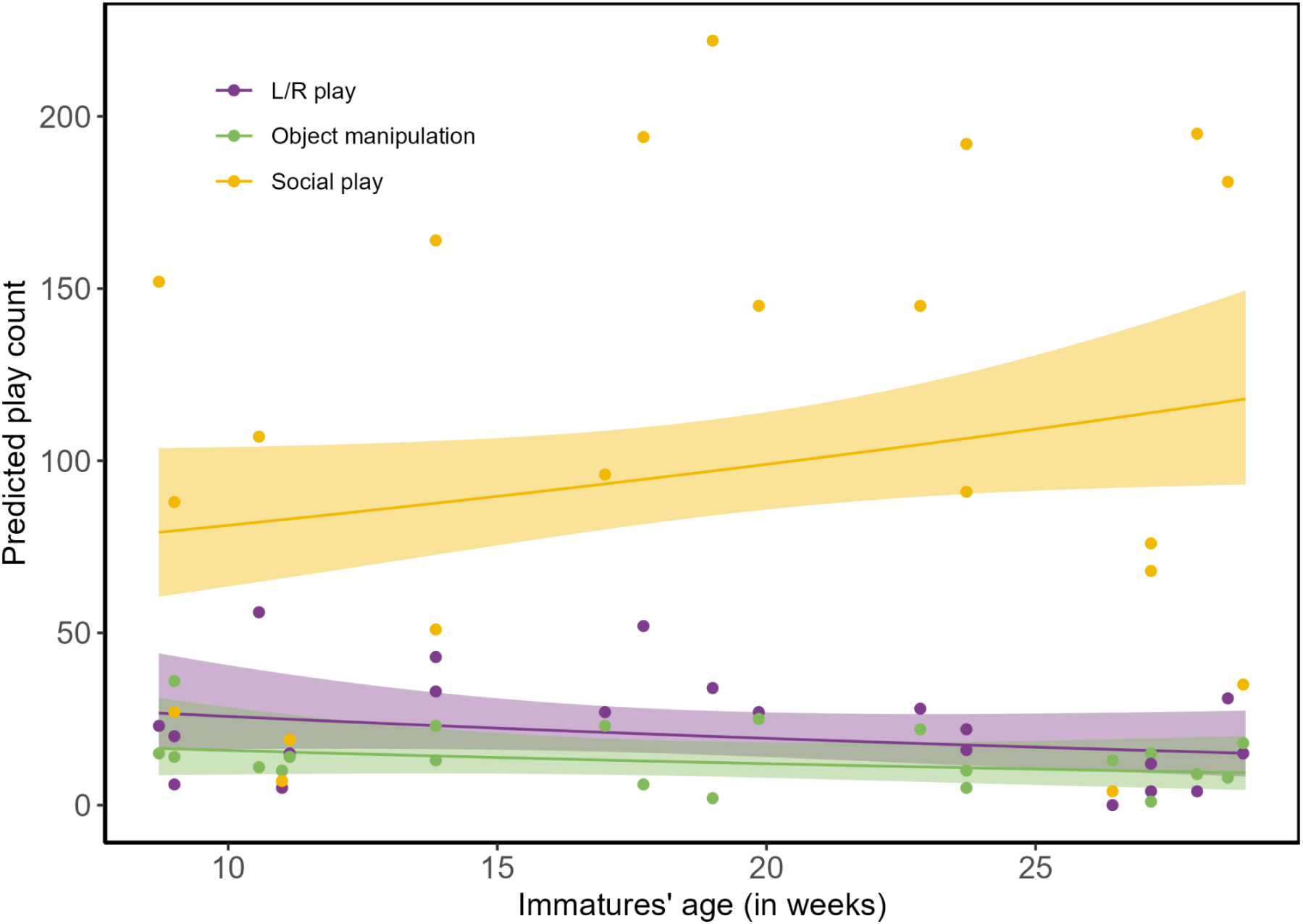
Predicted trajectories of the three play categories between 2 and 6 months old. Each point represents the frequency of a specific play category of a pair of twins from the same group (N_total_ = 63, N_twins_ = 3 pairs of twins, N_groups_ = 3). Shaded areas illustrate 95% confidence intervals. Raw data are shown as points. Points are raw counts of play elements in a single session without accounting for session duration and time out of sight.

**Table 1.**
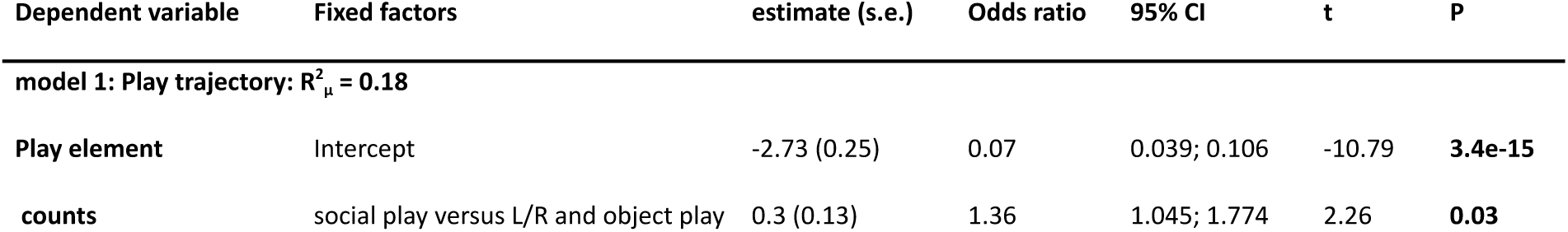
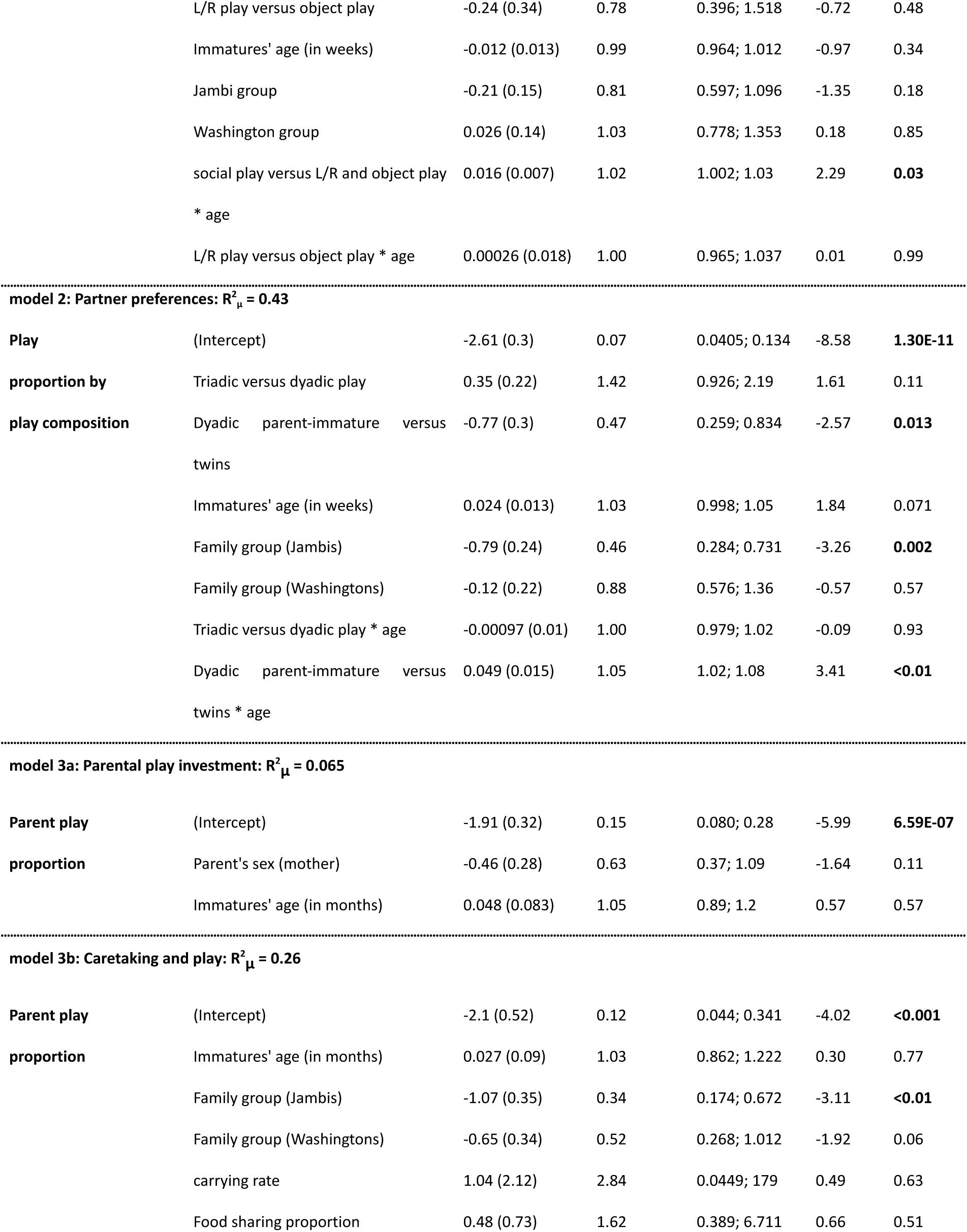
Summary of the GLM model 1 and Gamlss models 2 and 3. Bold values indicate significant predictors (p < 0.05)

### Part 2 : Play partners

Next our focus was on how much time an immature would spend playing with its twin (dyad), a parent (dyad) or her twin and a parent (triad) depending on age. The full model (AIC= –148.55) that included the fixed effects play composition, age of the immatures and group as well as the two-way interactions between age and play composition, explained the data significantly better than the null model (N_total_ = 63, N_individuals_ = 12, N_groups_ = 3; AIC = –126.79; likelihood ratio test: *X*^2^ (7) = 35.77, *p* < 0.0001).

We found a significant interaction between play partner composition (i.e.,Dyads_PARENT-IMMATURE,_ Dyads _TWINS_ and Triads_PARENT-TWINS_) and age of immatures. Planned contrasts showed that with immature age the odds of playing in a dyad composed of two immatures (Dyads _TWINS_) increased 5% faster compared to a dyad composed of a parent and an immature (Dyads_PARENT-IMMATURE_). No difference was found between the dyadic and triadic play in terms of rate of change with immature age (Table 1, model 2, Figure 3a).

**Figure 3.**
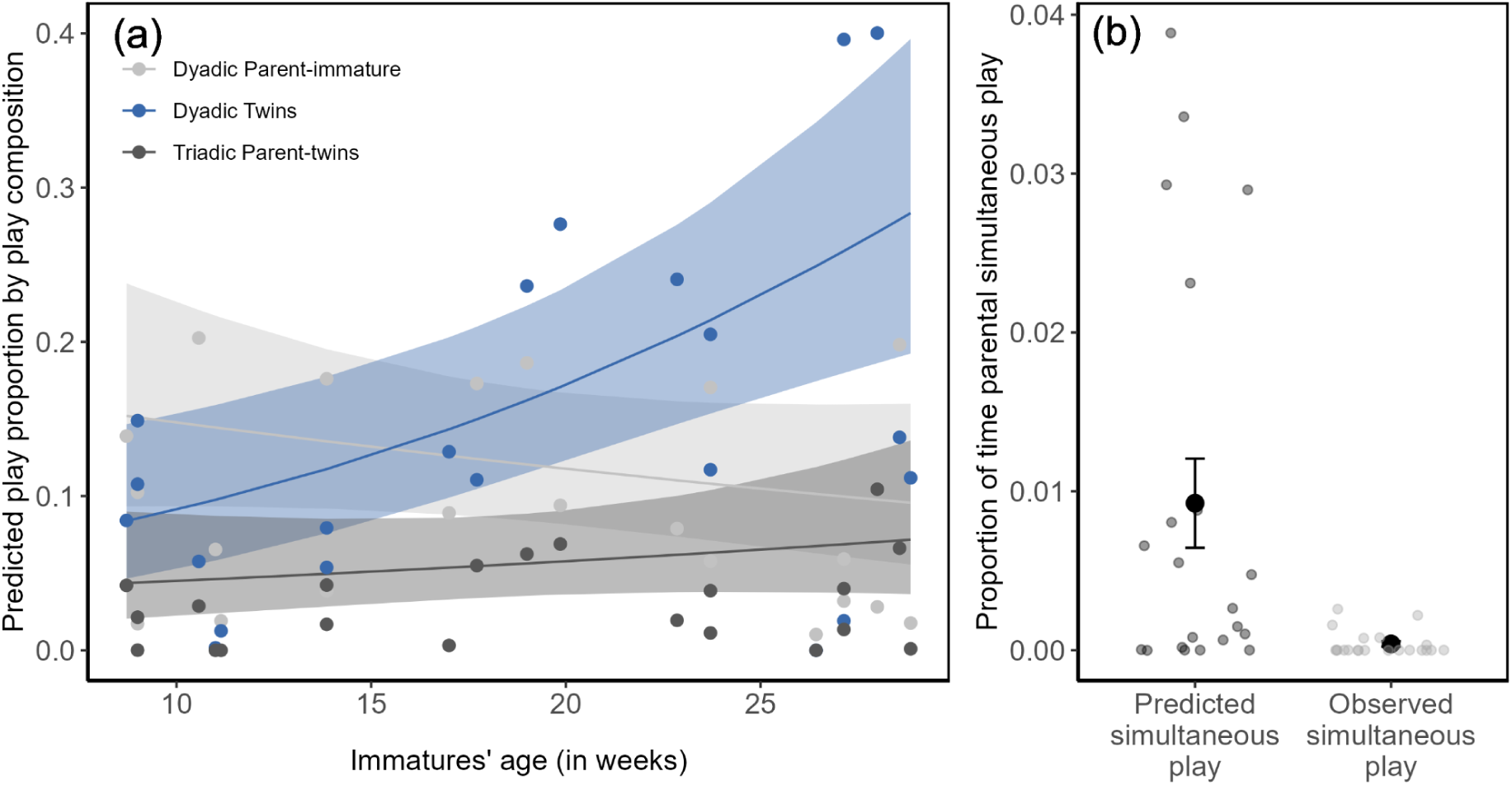
Partner preference and simultaneous parents play. (a) Predicted proportion of time spent playing in the different compositions over the 5 months of observation (model 2). Shaded areas illustrate 95% confidence intervals. Raw data are shown as points. A point represents the play proportion of a specific composition (i.e., Dyads_PARENT-IMMATURE,_ Dyads_TWINS_ and Triads_PARENT-TWINS_) in a single session (N_total_ = 63, N_individuals_ = 12, N_groups_ = 3). (b) Comparison between predicted and observed proportion of time individuals played simultaneously (N_total_ = 42, N_individuals_ = 6, N_groups_ = 3). Points represent the raw values of predicted (black) and observed (grey) simultaneous play per session. Error bars are based on raw data and show the standard error of the mean, the mean is represented with the opaque black dot.

On average, between 9 weeks old until 19 weeks old, an immature (independent of group identity) played in a dyad with one of their parents for 54% out of the total time spent playing in dyads. Between 19 weeks until 29 weeks old, out of the total time spent playing in a dyad, immatures played on average 29% with either one of their parents and the rest of the time spent playing, with their twin. This was driven by an increase of play between twins (Dyads _TWINS_) and not a decrease in parent-immature (Dyads_PARENT-IMMATURE_) play.

In addition we found group differences in the time spent playing overall (indifferent of the play composition) and post hoc tests identified the group “Jambi” as playing less compared to the group “Guapa” and “Washington” (see Figure S2, Table S4).

### Part 3: Parental investment and coordination

Finally, we investigated the difference between fathers and mothers in time spent playing with the immatures. The model (AIC= –51.8353) including the sex of the parent, the age of the immatures (in months) and group identity did not explain the time spent playing better than the null model (AIC= –53.0186). Contrary to our predictions, mothers and fathers played at equal amounts throughout the five months of observation (see Table 1, model 3a, Figure S3).

In addition, we examined the trade-offs between playing and other care-taking behaviours. The model (AIC = –55.4965) including carrying rate, food sharing proportion, age of the immatures in months and group identity did explain play involvement better than the null model (AIC= –53.0186), however this was driven by the group identity and not carrying or food sharing (see Table 1, model 3b).

Lastly, we wanted to quantify if and how much time parents played simultaneously either in triad or tetrad (mother-father-immature(s)), in two simultaneous dyads (father-immature and mother-immature) or with each other (father-mother). We examined all coded data and found a total of 6.43 seconds of overlap (over 1060 minutes of observation), equalling 0,0001 % of observation time. When comparing between the predicted and observed simultaneous play from the father and the mother we could show a significant difference between the proportions (z [20] = 3.41 ; p <0,001, r = –0.74), with lower observed values of simultaneous play than predicted values (Figure 3b). The parents were thus less likely to be playing at the same time than what would have been expected by chance.

## Discussion

In this study we investigated the ontogeny of play in three groups (N = 12) of cooperatively breeding common marmosets with immatures between the age of two to six months. We found that social play follows the expected trajectory, but not L/R and object play. Immatures’ partner preference changed depending on their age, with the twin becoming the more important play partner over time but adult play remaining similarly important (of the total play time when an immature played with a partner, this partner was in 54% of the time a parent during weeks 9 to 19 weeks old, and in 29% during weeks 19 to 29 weeks old). Lastly we did not find any differences in parental investment into play between mothers and fathers (model 3a) and indeed also did not find any support for trade-offs between other care-taking behaviours (such as food sharing or carrying) and play (model 3b). However, we found strong evidence that parents avoided playing at the same time.

**Part 1: Play trajectories**. The social play trajectory followed the expected pattern, with an increase of play frequency over the five months of observation. Since the peak of play is expected at the juvenile period, as in many other species where play has been investigated e.g., dogs (Pal, 2010), jungle babblers *Turdoides striatus* (Bond & Diamond, 2003; Gaston, 1977), gazelles *Gazella cuvieri* (Gomendio, 1988), and marmosets are considered juveniles between 3 months (Abbott et al., 2003) and 12 months old, it is unsurprising that the decrease in social play frequency was not yet observed in our observation period which ended at 6 months of age. However, the prediction we had made that all play categories would increase in frequency with age was invalidated. Both L/R play and object manipulation stayed stable throughout the five months of observation and were very low.

We confirmed that the different play categories are not equally distributed. Social play was much more common than L/R play and object manipulation throughout the five months of observation. This near absence of solitary play, at any age observed and the high and increasing level of social play could be explained with the idea that play reflects the ecology of a species. As cooperative breeders, marmosets possess strong social bonds and are heavily interdependent (Burkart et al., 2022; Koenig, 1998; Yamamoto & O.Box, 1997). Cooperation is key in marmosets amongst all family members and one hypothesis about the function of play is that it might enhance social skills. Our results combined with the evidence from other research on callitrichids support this notion especially since cooperative acts do not only happen among peers in callitrichid groups but all group members (Burkart et al., 2009). It makes sense then that the whole group participates in social play behaviours requiring a specific form of synchronisation with one’s partner, shared intentionality and coordination on the fly. On the other hand, common marmosets are exudate feeder–insectivores and frugivores. They have specific dental adaptations for tree gouging but none of their food sources (gum, fruits or insects) requires a technical manipulative component (Sussman and Kinzey 1984). This would explain the low frequency of object manipulation in marmosets, because they do not have to train any particular technical skills, but not the impact it has on its developmental trajectory. The hypothesis that object play in immatures might be a precursor to tool use and allow practice of technical skills required in adulthood has found some echoe in several species, such as in bonobos and chimpanzees (Koops et al., 2015). Moreover, in Cuvier’s gazelle *Gazella cuvieri,* a species where tool use is absent, Gomendio (1988) found that there was no effect of ontogeny on object play and it was generally very low, but still existent. Here, we show that even in non-tool users, a certain amount of object manipulation exists but the developmental trajectory does not follow the expected pattern. Despite the very low occurrence of object play at the very start, it would nonetheless be interesting to explore object manipulation in older marmosets to see if there is a decline in this behaviour later in development.

The results are even more surprising for solitary L/R play. As immatures become confident in their environment, we expected to see an increase in solitary locomotor play, but this did not happen. Perhaps, L/R play in marmosets only takes a social form and is incorporated into social play elements, such as play chase, reducing to a minimum solitary locomotor play elements. These results show that play is a heterogeneous behaviour, with different patterns for each category. One issue with coding L/R play, that could influence the results in general when studying L/R play, is that it can be difficult to disentangle from mere locomotion. If an individual is running, with no play face (the play face is seen more often in social play than solitary play (Ross et al., 2014) and alone, this individual is assumed to “just” move and not play, but a chance remains that the individual indeed was playing (for an in-depth discussion of reasons why solitary play is especially difficult to code, regarding the criteria established by Burghard, 2005 see section S2). We believe this to be a limitation when studying solitary play in general.

**Part 2: Play Partners.** We then investigated the immature play partner preferences. As predicted, twin play increased with age, but the decrease expected in parent-immature play was not observed, perhaps this happens later in development during the juvenile period. Data on a larger sample size with older immatures would therefore be required to fully capture the development of partner preference over time in marmosets.

Notably, parents were important partners during playful interactions throughout the observation period, despite the constant availability of a same-aged partner (the twin). This contrasts with what has been observed in squirrel monkeys (Biben, 1998), for which the low involvement of the mothers has been attributed to the presence of peers, with the underlying idea that play would cost mothers energy and time that they might not be able to afford. Similar patterns of a strong preference of immatures to play with individuals as close as possible in age, at any age observed, have been reported in tamarins (Kostan & Snowdon, 2002; Epple & Katz, 1980), meerkats *Suricata suricatta* (Sharpe, 2005) and western lowland gorillas *Gorilla gorilla* (Maestripieri & Ross, 2004). One possible explanation for such high levels of parental involvement in play could be the captive housing condition reducing the energy demands on parents and thus allowing them to play more. What speaks against this argument is that studies conducted on other species in captivity have not found that parents are more involved in play e.g., hyaenas *Crocuta crocuta* (Drea et al., 1996), or western lowland gorillas (Maestripieri & Ross, 2004). It is important to note that the groups included in the sample for this study did not include any helpers thus very likely the amount of mixed-age play is underestimated, as the majority of studies in callitrichids show more play by juvenile and subadult helpers than breeders (Cleveland & Snowdon, 1984; De Oliveira et al., 2003; Stevenson & Poole, 1982). Interestingly, our prediction that polyadic play should be less commonly observed than dyadic play was invalidated. Triadic play was as prevalent as dyadic play throughout the five months of observation. This result reinforces the idea that marmosets can easily coordinate play, even in a triad, without the risk of escalating into a real fight.

Lastly, group identity seemed to have had an important impact on how much an individual played, for the immatures as well as for the parents. All individuals in the Jambi group played less than the individuals in the Guapa group, raising the question as to whether different amounts of play are part of “group personalities” in playfulness. Previous research on marmosets has shown group-level similarities (i.e., “group personalities”) on traits such as boldness and exploration (Koski & Burkart, 2015). These traits were better explained by social mechanisms (social proximity within the group) than genetic relatedness between individuals. To strengthen the hypothesis that playfulness is a group personality trait and to understand how flexible this trait is, the study of more groups would be required as well as the monitoring of those group members over time to confirm consistency and the changes in playfulness or not when individuals change group to form a new family. A recent study investigated the social function of play signals in the same groups of common marmosets. Results showed that marmosets use body posture as a multi-functional signal to initiate and prolong their play interaction, as well as allowing for more intense play (Adriaense et al., unpublished data) while acknowledging group differences. These results further highlight the potential for differing play communication styles between marmoset groups.

**Part 3: Parental investment and coordination.** Finally, we compared investment in play by mothers and fathers, and investigated if parental play was correlated with infant care contributions i.e., food sharing and carrying. There was no significant difference between mothers and fathers with regard to the time they spent playing with their offspring. This result hence contradicted our prediction by showing that in marmosets, both parents play equally. This is surprising given that females are under higher energetical demand compared to males (Erb & Porter 2017). In the species where parents do get involved in playful interactions, especially at a young age, these playful interactions are often exhibited with the primary caregiver i.e., mothers in chimpanzees Pan troglodytes (Pellegrini & Smith, 2005) or orangutans Pongo spp (Kunz et al., 2024). In species where males take an active role in immatures caretaking, they also show an important involvement in playful interactions, sometimes more than the mothers: In Javan gibbons Hylobates moloch, fathers play more and groom the infants more than mothers do (Yi et al., 2023). In humans, who are cooperative breeders as the common marmosets, even if there is no final consensus on the comparison of father-child/mother-child play frequency (Amodia-Bidakowska et al., 2020) it is clear that both parents are very involved in playful interactions (Smith & StGeorge, 2023). Some studies even found play to be equally distributed amongst parents (Laflamme et al., 2002). In addition, Amodia-Bidakowska et al. (2020) also found that investment in play behaviours was dependent on parental employment status, thus dependent on the time available to allocate to play behaviour.

We expected that energetic constraints might factor into the decision of how much time is spent playing. Our result is therefore surprising as we expected the parent with more energetic constraints, i.e., the mothers, to play less compared to fathers. Since there was no impact of the parents’ sex on play investment, we checked whether a difference would arise at the individual level depending on how those individuals were involved in immature care taking. But neither carrying nor food sharing were predictive of how much an individual would play with her offspring. None of our competing hypotheses were thus validated, because there was neither a positive nor a negative correlation between caretaking behaviours and play. This result hints at the idea that play might be a special form of parent-offspring interaction or that our sample simply lacks statistical power to show a potential correlation between these behaviours.

Most intriguingly, there were almost no occasions in 1060 minutes of observation time, where both parents played simultaneously, it was always either the father or the mother with the immatures. The fact that they alternated who was playing with the immatures could have two explanations. The first is that adults do not often play together in general, in all species where we know play exists (Palagi, 2023; Pellis & Iwaniuk, 2000), likely in part to avoid the risk of escalation into real fights. In highly socially tolerant callitrichids this explanation is unlikely as in bigger groups with only adults play between adults still occurs (Norscia & Palagi 2011, personal observation in wild common marmosets by J.M.B.). The second explanation is linked to the potential threat individuals face while playing. Play can be costly for small prey animals because vigilance and play are hardly possible at the same time (Beauchamp 2015; Biben, 1998; Harcourt, 1991; Hausfater, 1976), since both require the attention of the player. If both parents would play at the same time, that would make immatures very vulnerable to predators, especially in small groups such as the ones included in this sample. This alternation between parents appears to be similar to the coordination capacity that marmosets show in a different context, namely when coordinating being vigilant and feeding head-down, in a situation where both of these behaviours are mutually exclusive (Brügger et al., 2023; Phaniraj et al., 2023). In a study on golden lion tamarins, De Oliveira et al. (2003) showed that adult breeders increase their vigilance when immatures are playing. This would hint again at the fact that adults do avoid playing together to make sure at least one individual can be vigilant in this dangerous context.

This study shows the clear preference of cooperatively breeding common marmosets for social play that increases with age over months two to six of their development. More importantly it highlights the large and similar involvement of both parents in play even though same aged twins and thus a peer to play would always be available. We do not find any support for trade-offs between other care-giving behaviours and play. Besides showing general patterns of immature play in this species we provide a detailed illustrated ethogram of the observed play behaviours with accompanying supplementary video material of all coded play behaviours encouraging future research on this highly dynamic behaviour in this species.

## Authors’ contributions

**A.M.G.:** Conceptualization, Methodology, Formal analysis, Investigation, Data Curation, Writing - Original Draft, Writing - Review & Editing, Visualization; **J.M.B.:** Conceptualization, Methodology, Writing - Original Draft, Writing - Review & Editing, Resources, Funding acquisition, Supervision; **R.K.B.:** Conceptualization, Methodology, Formal analysis, Investigation, Writing - Original Draft, Writing - Review & Editing, Supervision.

## Data availability

All relevant data are available on OSF at https://osf.io/wamnr/?view_only=002d0054aec64e0ab802a1637c8e70bc

## Acknowledgements

We thank Sandro Sehner for generously sharing his food sharing data, Melanie Meyer for inter-observer reliability coding and Fabian Hervas Peters for help with programming of the data extraction pipeline. We are grateful to Hidir Sengül and Dominique Ziegler for animal care-taking and support during data collection. We thank Erik Willems for guidance with statistical modelling and Jessie Adriaense as well as Anouk Manzanell for discussions that were instrumental in shaping the play ethogram.

