## Supplementary materials for "The ontogeny of play in a highly cooperative monkey, the common marmoset"

**Table S1.** Group composition

**Table S2.** Coding protocol for each of the play elements with focal and receiver of each element

**Table S3.** Inter-observer reliability for all the play elements coded.

**Table S4.** Estimated marginal means pairwise comparison between groups based on model 2

**Section S1**. Supplementary methods

**Table S5.** All interval ceilings (when maximum play duration is reached) for the three possible dyads in all sessions coded for the three groups

**Figure S1.** Example of how the interval changes play duration

**Section S2.** Coding solitary play

**Figure S2.** Predicted overall difference in amount of play per group based on model 2

**Figure S3.** Proportion of time fathers and mothers spent playing overall over the five months of observation

**Section S3.** Supplementary videos

Subtitles in the videos do not indicate the duration and start/end of a behaviour but only the presence of this behaviour. All videos include a real time and slowed down (0.6 or 0.4 speed) version of the respective elements and may also include other play elements than the one mentioned in the description.

**Video S1.** Social play, **wrestle**

**Video S2**. Social play, **grip**

**Video S3a.** Social play, **pull** (tufts)

**Video S3b.** Social play, **pull** (tail)

**Video S4.** Social play, **touch**

**Video S5.** Social play, **pounce**

**Video S6.** Social play, **chase**

**Video S7.** Social play, **catch**

**Video S8.** Social play, **supine**

**Video S9.** Object play, **manipulation**

**Video S10.** L/R play, **roll**

**Video S11.** L/R play, **somersault**

**Video S12.** L/R play, **acrobatic play**

**Video S13.** L/R play, **pounce on hammock**

**Video S14.** L/R play, **single stalk**

**Table S1.** Group composition.

| **Group** | Group composition | Individuals | Sex | Status | Birth Date | Age of inf/juv during recording period | Data collection period | Food sharing |
|---|---|---|---|---|---|---|---|---|
| **Guapa** | 2 b, 2 im | Guapa | f | breeder | 4 May 2018 | 2–6 months | May–Sep. 2021<br>*2 series/month* | May–Sep. 2021<br>*Twice a week* |
| | | Ninja | m | breeder | 12 Apr. 2015 | | | |
| | | Guarana | f | immature | 5 Mar. 2021 | | | |
| | | Gimli | m | immature | 5 Mar. 2021 | | | |
| **Jambi** | 2 b, 2 im | Jambi | f | breeder | 10 Nov. 2015 | 2–6 months | Sep–Jan 2021/22<br>*2 series/month* | Sep–Jan 2021/22<br>*Twice a week* |
| | | Werewolf | m | breeder | 10 May 2018 | | | |
| | | Jafar | m | immature | 7 Jul. 2021 | | | |
| | | Jagger | m | immature | 7 Jul. 2021 | | | |
| **Washington** | 2 b, 2 im | Washington | f | breeder | 30 Aug. 2013 | 2–6 months | Oct–Feb 2021/22<br>*2 series/month* | Oct–Feb 2021/22<br>*Twice a week* |
| | | Lotus | m | breeder | 3 Jul. 2012 | | | |
| | | Wall-e | f | immature | 12 Aug. 2021 | | | |
| | | Woody | m | immature | 12 Aug. 2021 | | | |

b = breeder, im = immature, 1 series = 1 video in the morning and 1 video in the afternoon in the same week

**Table S2.** Coding protocol for each of the play elements with focal and receiver of each element.

| Play component | Definition | Focal | Receiver |
|---|---|---|---|
| **Wrestle** | Starts when two individuals make physical contact (1st frame of contact). Initiator is the one making 1st contact exactly after a pull, grip or pounce (keep initiator of preceding component). | The individual making the first contact | The individual receiving the contact |

| | | | |
|---|---|---|---|
| | Ends when one of the players stops wrestling: 1) one subject jumps out of the wrestle (1st frame where one of the player moves a limb away from the other player) OR 2) one subject starts looking elsewhere (gaze away, vigilance) and stops interacting with the partner. It can also be that both of them stop at the same time. | | |
| **Grip** | Unilateral grasp from behind or side of the play partner. Starts at the first frame where both hands are around the play partner, as long as contact lasts or as long as it is unilateral i.e., can shift to wrestle when the individual gripped turns around to face the attacker.<br>End from the grasp is released. | The individual gripping | The individual gripped |
| **Pull** | Pull starts when first contact happens between the individual that is pulling and the one that has e.g., her tail pulled. Does not count when part of wrestle.<br>Ends when the contact between the players had ended or the pull had shifted to another component. | The individual pulling | The individual pulled |
| **Touch** | Touch coded when the one individual touches, pushes or slaps a playmate with the hand (or 2 hands). The one touching should be looking at the receiver. Does not count when part of wrestle. Do not code touch if preceding wrestling, gripping and pulling. | The individual performing the touch/slap/push | The individual touched/slapped/pushed |
| **Pounce** | An individual pounces with a circular arc trajectory in an arched back position (cat pose) while landing. At least one of the limbs touches the play partner (any part of the playmate doesn't have to be fully on the body). The landing should be with the 4 limbs at the same time. | The individual pouncing | The individual being pounced on |

| | | | |
|---|---|---|---|
| **Chase** | Starts at 1st frame when both individuals have initiated their first movement: i.e., 1st frame where the last player to start running ("the chaser") has a front limb that does not touch the substrate anymore. The individuals should be running one behind the other most of the time. Parallel run can happen during chase but it should be following or followed by a clear run directed at the chasee. Any pounce on hammock performed during the chase is not counted since the chase has priority over solitary events.<br>Ends when the chaser stops chasing the other/chasee (1st frame where all four limbs are touching the substrate) or manages to catch up with the chasee (1st frame of contact). | Chaser | Chasee |
| **Catch** | Starts at 1st frame with one limb not touching the substrate as the (jerky) movement is initiated. Happens on big branches and grid. 2 individuals do it at the same time with one moving backward and the other trying to catch/touch her.<br>Ends when at least one individual has disengaged from the catching-escaping game. | The first individual trying to catch the other one in the stalk | The first individual that is avoiding the catch by stalking in return |
| **Supine** | An individual is lying on the back or on the side. 2 legs at least should be lifted.<br>If the receiver is not visible or if gaze is pointing towards 2 individuals, code UK (unknown).<br>The hands/feet should not be around a branch. | The individual doing the supine | The individual gazed at |
| **Manipulation** | Starts at the 1st frame where an individual grasps, takes, mouth or gnaws a fixed or free object without using it for any specific goal (non-edible objects, doesn't include camera nor other individuals nor branches).<br>The manipulation is therefore restricted to the | The individual manipulating | No |

| | | | |
|---|---|---|---|
| | hammock, ropes, strings and any inedible objects that are in the enclosure. Manipulation should be done with the attention of the manipulator, meaning that the gaze should be turned towards the object manipulated.<br>Ends when an individual either stops looking at the object or releases the object from her grip. | | |
| **Roll** | Voluntary' side roll: do not count rolls on the hammock that happen because an individual is falling at the bottom of the hammock. | The individual rolling | No |
| **Somersault** | rolling over the head, can be a bit on the side, but the head should come first in any instances of a somersault, in contrary to a roll. | The individual doing the somersault | No |
| **Acrobatic play** | 1st frame with at least one limb touching the unstable substrate of interest (rope, hammock, twig) and other limbs not in contact anymore with a stable substrate (grid, branch). At least one hand/feet should hang loose.e.g., swing around rope, hang from twig or hammock.<br>Exclude if 1) if an individual uses a rope to climb on the hammock (individual moving on every frame), the use of the rope is a means to get to somewhere else and the rope is not the goal per se.<br>Ends when the individual is touching again a stable substrate or jumps away. | The individual doing acrobatic postures | No |
| **Pounce on hammock** | 1st frame where at least one of the front limbs is not on the initial substrate as the jump is initiated toward the hammock. The landing should be with the 4 limbs at the same time, hence when the landing is with only the 2 front limbs first it is not considered as a pounce (see examples in the components folder). The pounce has a circular arc trajectory and the individual has an arched back | The individual pouncing | The individual(s) present on the hammock |

| | | | |
|---|---|---|---|
| | position (cat pose) while landing. Do not count as receivers, individuals that are on the ropes of the hammock. | | |
| **Single stalk** | Starts at 1st frame with one limb not touching the substrate (branch) as the (jerky) movement is initiated. Happens on big branches and grid mainly. Is combined with gaze/mutual gaze towards another individual. Ends when the individual is completely static again. | The individual doing the stalk | The individual gazed at |
| **UK** | Any play element that is not distinguishable because too far away or that is not defined in the ethogram, for example : -bites when not combined with any other component -unilateral wrestles -manipulation of own or others tail | The individual performing a play element | The individual receiving the play element |
| **Out of sight (oos)** | Any instances where an individual is not visible at all anymore. | The individual out of sight | No |

**Table S3.** Inter-observer reliability for all the play elements coded.

| Play element | ICC | duration(D)/ occurrence(O) |
|---|---|---|
| chase | 0.97 | D |
| wrestle | 0.99 | D |
| pull | 0.93 | D |
| stalk | 0.93 | D |
| catch | 0.89 | D |
| grip | 0.93 | D |
| supine | 0.86 | O |
| touch | 0.81 | O |

| | | |
|---|---|---|
| pounce | 0.96 | O |
| pounce on hammock | 0.99 | O |
| manipulation | 0.98 | D |
| acrobatic play | 0.38 | D |
| somersault | excluded | O |
| roll | excluded | O |

**Table S4.** Estimated marginal means pairwise comparison based on model 2. The pairwise comparison occurs between the three family groups, indifferent to the play composition . Bold values indicate $p < 0.05$.

**model 2 : play proportion ~ play composition* immatures' age ( in weeks) + family group**

**Pairwise comparison between the three family groups, indifferent of the play composition**

| **contrast** | **estimate (SE)** | **95 % CI** | **z-ratio** | **p-value** |
|---|---|---|---|---|
| Guapa versus Jambi group | 0.785 (0.241) | 0.220; 1.35 | 3.258 | **0.0032** |
| Guapa versus Washington group | 0.124 (0.218) | -0.388; 0.636 | 0.569 | 0.837 |
| Jambi versus Washington group | -0.661 (0.242) | -1.228; -0.0933 | -2.729 | **0.0175** |

**Section S1.** Supplementary methods.

We varied intervals from 1 second to 60 seconds to see if the time spent playing between the same dyad, not joined by another individual, would change depending on the length of the chosen interval. The process consisted of first going through the coded play bout with an interval of 1 second and link all the play elements that were closer to another than 1 second if they were between the same dyad. All moments where partners changed or one player joined were therefore excluded from the time spent playing for the dyad in question. In other words, if two play elements were less than a certain number of seconds apart from each other and performed by the same pair of players, these were treated continuously as the behaviour 'play'. We repeated this process increasing the interval duration of one second each time. We found that from a certain interval on, the play duration per dyad was reaching a ceiling point. We decided to take the middle point between 0 and the mean ceiling interval length as our interval duration, which corresponded to 10 seconds.

**Table S5.** All interval ceilings (when maximum play duration is reached) for the three possible dyads in all sessions coded for the three groups.

| **Group** | **Immatures' age (months)** | **Ceiling Dyads** $_{\text{TWINS}}$ | **Ceiling Dyads** $_{\text{FATHER-IMMATURE}}$ | **Ceiling Dyads** $_{\text{MOTHER-IMMATURE}}$ |
|---|---|---|---|---|
| Guapa | 2 | 29 | 12 | 25 |
| Guapa | 2 | 9 | No play | 35 |
| Guapa | 3 | 11 | 13 | 42 |
| Guapa | 4 | 17 | 14 | 12 |
| Guapa | 5 | 20 | 16 | 12 |
| Guapa | 6 | No play | No play | 1 |
| Guapa | 6 | 9 | 48 | 14 |
| Washington | 2 | 28 | 13 | 11 |

| | | | | | |
|---|---|---|---|---|---|
| Washington | 2 | 15 | 1 | 1 | |
| Washington | 3 | 24 | 24 | 30 | |
| Washington | 4 | 23 | 19 | 23 | |
| Washington | 5 | 16 | 6 | 10 | |
| Washington | 6 | 19 | 1 | 22 | |
| Washington | 6 | 15 | 9 | No play | |
| Jambi | 2 | 36 | 1 | 2 | |
| Jambi | 2 | 38 | 12 | 1 | |
| Jambi | 3 | 57 | 18 | 36 | |
| Jambi | 4 | 21 | 14 | 2 | |
| Jambi | 5 | 32 | 11 | 28 | |
| Jambi | 6 | 50 | 12 | No play | |
| Jambi | 6 | 31 | No play | 11 | |
| **Mean** | | 25,00 | 13,56 | 16,74 | **18,43** |

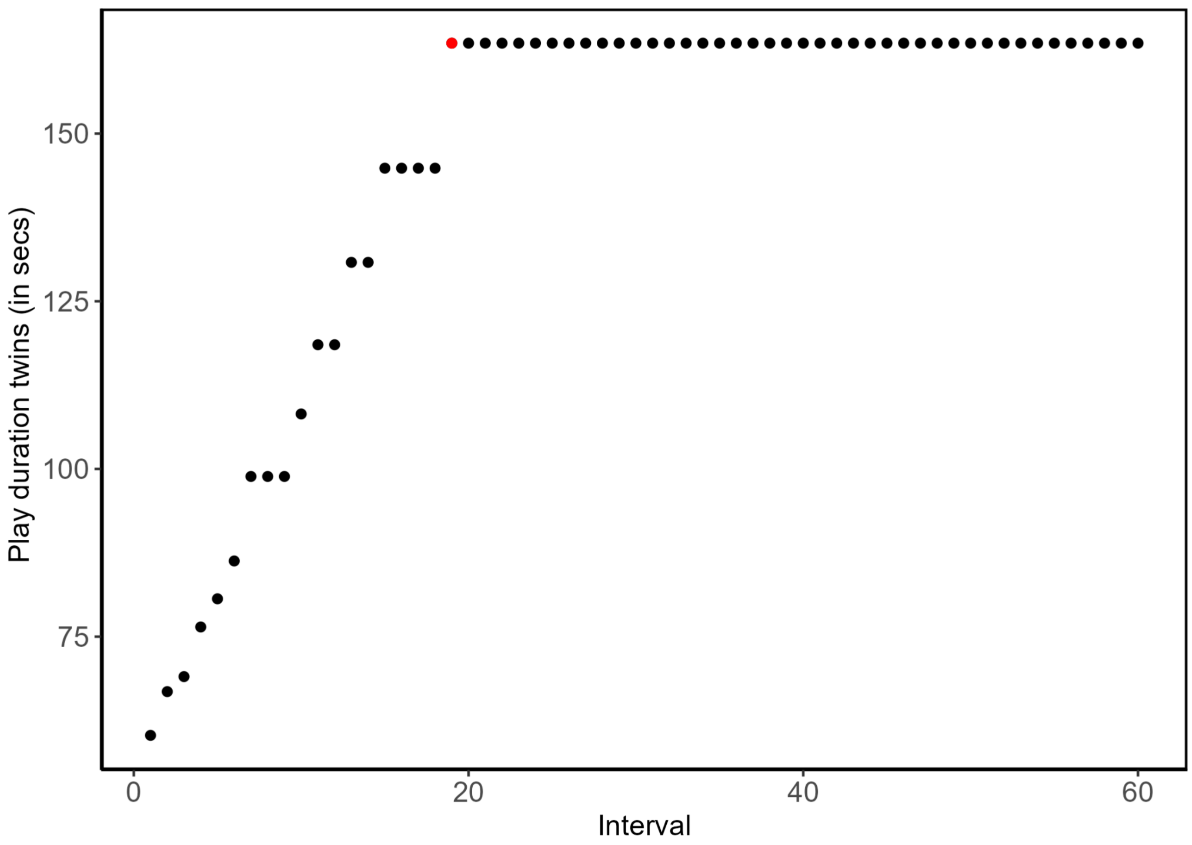


**Figure S1.** Example of how the interval changes play duration. Depending on the interval length (varying from 1 to 60 seconds), play duration also varies, but reaches a ceiling point nonetheless during the 60 seconds. Here, the ceiling point is reached when the interval is 19 seconds (point shown in red), meaning that the play elements during a play bout between twins in this particular session were always at a maximal distance of 19 seconds apart.

**Section S2.** Coding solitary play

Running can fulfil all criteria of play (Burghard, 2005; see Methods, section 'Data coding and preparation' for a full description of the 5 criteria), but it is context-dependent and delicate to categorise especially because of the first and third criteria. The first criterion mentions that the behaviour has to be not fully functional by including elements that do not contribute to immediate survival. In the case of running or pouncing, it is very difficult to know or infer the reason why an individual might have initiated these behaviours. It could be because they heard a noise that humans can't hear on the video, a sudden desire to be closer to her twin or group mate, or it could be play. Moreover, it is difficult to differentiate, for fast arboreal primates such as marmosets especially, between a 'normal' run (i.e. just locomotion) without added elements and a run that is more acrobatic and less functional or efficient. The third criterion mentions that the behaviour has to be either incomplete, exaggerated, awkward, precocious, or modified. Again similar questions as to the first criterion apply. Due to these reasons our coding for L/R play was very conservative to avoid false positives, but therefore could underestimate the true frequency of L/R play.

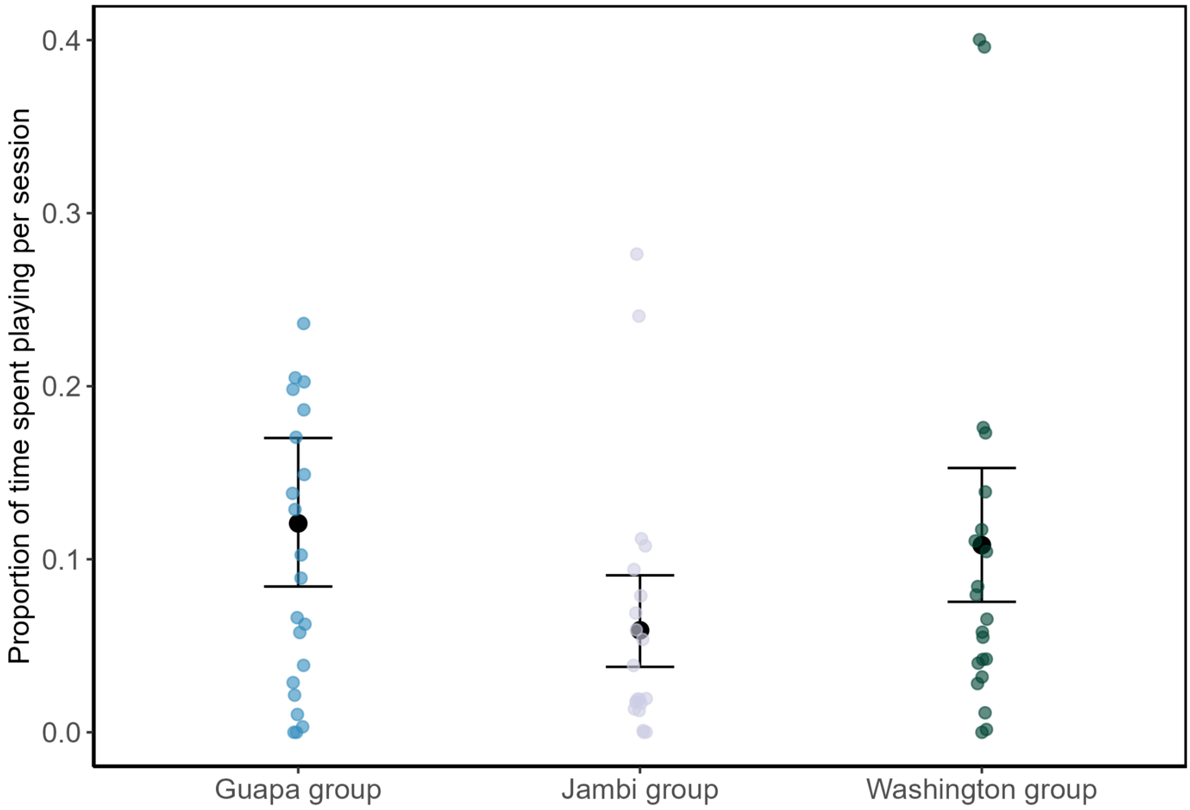


**Figure S2.** Predicted overall difference in amount of play per group. Each point represents the proportion of play between a dyad (indifferent of the dyad type) and error bars show 95% confidence intervals obtained from the model ($N_{total}$ = 63, $N_{individuals}$ = 12, $N_{groups}$ = 3). Raw data are shown as points.

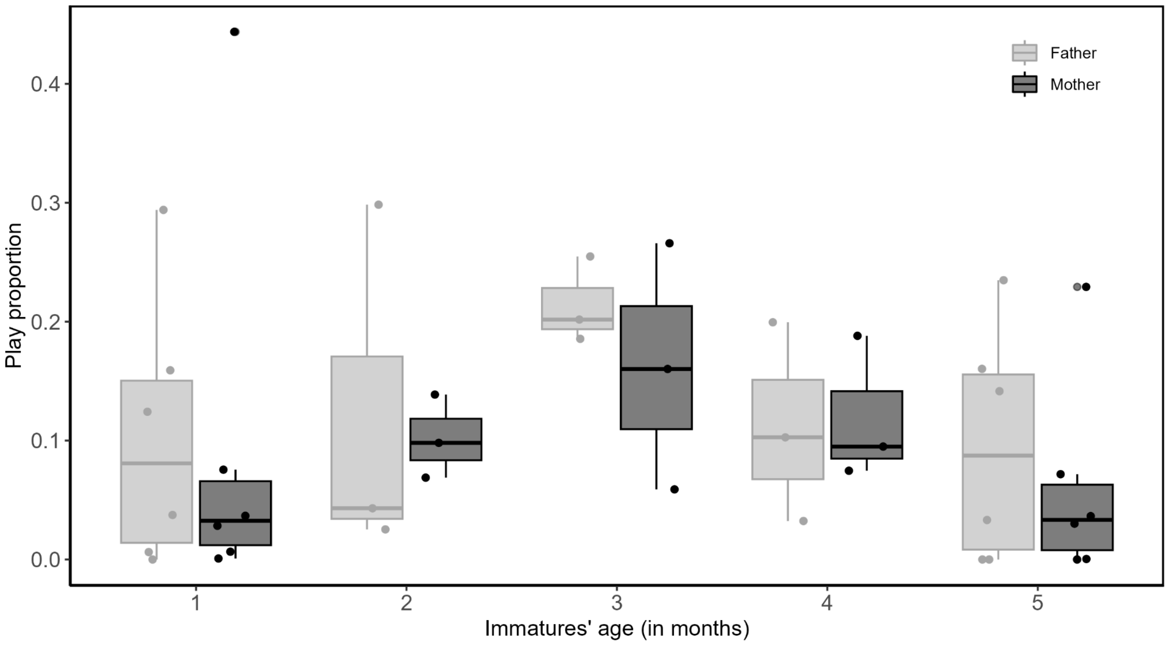


Figure S3. Proportion of time fathers and mothers spent playing overall over the five months of observation. Box plots and points are based on raw values. Boxes represent the 25th and 75th percentiles (ends) and the median (centre line); whiskers represent the first quartile − 1.5 × the interquartile range and the third quartile + 1.5 × the interquartile range and all outliers beyond these whiskers are shown.
